# Plant immunity activated and suppressed by a conserved effector protein family in mirid bug *Riptortus pedestris*

**DOI:** 10.1101/2023.08.22.554304

**Authors:** Jiangxuan Zhou, Zhiyuan Yin, Danyu Shen, Yumei Dong, Yuxia Yang, Qingsong Zhang, Yurong Ma, Yong Pei, Wangshan Lu, Yancong Zhang, Gan Ai, Donglei Yang, Yuanchao Wang, Daolong Dou, Ai Xia

## Abstract

*Riptortus pedestris* (Fabricius) a major soybean pest migrates into soybean fields during pod filling stage resulting in a leaf and stem staygreen while pods without beans syndrome. Given the agricultural importance of this species and the lack of characterized HAMP from piercing-sucking insects we performed a large scale of screening by expression of 87 *R. pedestris* salivary proteins with signal peptides in *Nicotiana benthamiana* obtaining a candidate HAMP RPH1. RPH1 activated a series of PTI responses including ROS burst upregulation of defense marker genes such as PR1 WRKY7 WRKY8 Acre31 and CYP71D20 MAPK activation and biosynthesis of phytohormones in plants. RPH1 significantly enhances soybean resistance against *R. pedestris* feeding. PRR coreceptors BAK1 and SOBIR1 were required for RPH1-induced PTI responses. Remarkably RPH1 homologs were widely distributed in herbivorous insects and majority of homologs from selected species induced cell death or ROS. Thus our results demonstrated that RPH1 is a conserved HAMP within chewing and piercing-sucking insects. We also discovered that *R. pedestris* evolved four paralogs to overcome the plant immunity triggered by RPH1. This study filled a major gap of HAMP identification from piercing-sucking insect and also deciphered a novel evasion strategy of plant immunity exploited by herbivorous insects.

**One sentence summary:** *Riptortus pedestris* RPH1, a conserved HAMP in herbivores, activates a variety of PTI responses in plants. To couterdefense, *R. pedestris* evolved four paralogs to suppress RPH1-induced PTI responses.

The author(s) responsible for distribution of materials integral to the findings presented in this article in accordance with the policy described in the Instructions for Authors (https://academic.oup.com/plcell/pages/General-Instructions) is: Ai Xia.

## Introduction

*Riptortus pedestris* (Fabricius) is widely distributed in East Asia and an omnivorous pest with a broad range of host plants, but mainly infesting soybean (*Glycine max*) (Jung and Lee, 2018). It has caused tremendous yield losses for soybean growers since it withdraws nutriments from soybean pods using its needle-like mouthpart, resulting in a staygreen syndrome, characterized by delayed leaf and stem senescence, and pods without beans (Li et al., 2019; Jin et al., 2022). Like other piercing-sucking herbivores, *R. pedestris* also delivers a repertoire of effectors into soybean to promote pest infestation (Wei et al., 2023; Dong et al., 2022). For instance, *R. pedestris* effector RP2155 impairs soybean defense by suppressing the jasmonic acid (JA) and salicylic acid (SA) pathways (Huang et al., 2023). Silencing of RP614 significantly attenuated staygreen syndrome (Shan et al., 2023). However, how soybean recognizes salivary proteins released by *R. pedestris* and subsequently activates a broad spectrum of immunity is still unclear.

Plants’ recognition of molecular cues from herbivores to mount successful resistance is the key for their survival. Analogous to microbe-associated molecular patterns (MAMPs), herbivore-associated molecular patterns (HAMPs) are defined as molecules derived from herbivores which bind to pattern recognition receptors (PRRs) to trigger pattern-triggered immunity (PTI) (Mithofer and Boland, 2008). Phytophagous pests are generally grouped as chewing and piercing-sucking insects (Bonaventure G, 2012). Chewing herbivores release a vast amount of saliva into plants when engulfing plant tissues. By collecting secreted saliva components, several HAMPs have been isolated including fatty acid conjugates (FACs), caeliferins, inceptins, and salivary enzymes such as b-glucosidase and glucose oxidase (GOX) (Acevedo et al., 2015). Volicitin was the first characterized HAMP from fall armyworm *(Spodoptera frugiperda)* (Alborn et al., 1997). Radiolabeled volicitin was reported to bind to a plasma membrane protein from maize (Truitt et al., 2004). *S. frugiperda* inceptins, proteolytic fragments of plant chloroplastic ATP synthase gamma subunit (cATPC), represent the best studied HAMPs so far (Schmelz et al., 2006; Schmelz et al., 2007). A leucine-rich repeat receptor-like protein (RLP), termed INR, confers inceptin-induced signaling and defense outputs (Steinbrenner et al., 2020).

In contrast to chewing herbivores, piercing-sucking insects secrete only small amounts of saliva into hosts during feeding, rendering it difficult to purify HAMPs from insect oral secretions (Bonaventure G, 2012). By transient expression of salivary proteins predicted from salivary gland transcriptomes or proteomes in plants, a few proteins including mucin-like salivary protein (NlMLP), LsPDI1, NlVgN, NlDNAJB9 and CathB3 have been identified to elicit plant defenses (Snoeck et al., 2022; Gao et al., 2023; Fu et al., 2020); However, these salivary proteins activate plant immunity within cells. For instance, the brown planthopper (BPH) NlMLP triggered immune responses in the cytoplasm of rice (Shangguan et al., 2018). *Myzus persicae* salivary protease CathB3 binds to cytoplasmic kinase ENHANCED DISEASE RESISTANCE 1-like (EDR1-like) and activates plant defense in phloem cells (Guo et al., 2020). Thus, it is still elusive whether HAMPs exist in piercing-sucking insects.

Host-adapted plant pathogens have evolved various strategies to evade plant immunity including extracellular strategies to avoid MAMP recognition, modulation of plant immune signaling, and disarming plant immune outputs (Wang et al, 2022). Similar to plant pathogens, phytophagous insects also inject effectors into host cells to suppress plant immunity triggered by PRR perception (Erb and Reymond, 2019). For example, *Laodelphax striatellus* and *Apolygus lucorum* deploy vitellogenin and Al106 to reduce reactive oxygen species accumulation for promoting insect performance (Ji et al., 2021; Dong et al., 2020). *Bemisia tabaci* BtE3 and *Helicoverpa armigera* HAS1 block JA signaling to dampen plant defenses (Chen et al., 2023; Peng et al., 2023). An extracellular strategy to prevent HAMP perception was also exploited by specialist caterpillar *Anticarsia gemmatalis* which processed inceptin-related peptides into a biologically inactive form, namely Vu-In2A, to avoid plant receptor perception (Schmelz et al., 2007; Schmelz et al., 2012). Despite these advances, our understanding of evasion of plant immunity by herbivores is still limited in comparison with the well-studied adaptation mechanisms of plant pathogens.

Here, a systematic screening was performed to search for candidate HAMPs from *R. pedestris*. The search yielded a HAMP which was termed RPH1 (*Riptortus pedestris* HAMP 1). Induction of plant defenses by RPH1 required PRR co-receptors BAK1 and SOBIR1. Furthermore, RPH1 was found to be a conserved HAMP existing in diverse chewing and piercing-sucking herbivores. Finally, we deciphered that *R. pedestris* has evolved paralogs of RPH1 to evade plant immunity triggered by RPH1.

## Results

### RPH1 from *Riptortus pedestris* triggered PTI responses in *N. benthamiana*

Salivary proteins delivered into soybean leaves during *R. pedestris* infestation were characterized using liquid chromatograph mass spectrometry (LC-MS). Using the soybean and *R. pedestris* genomes as references (Huang et al., 2021), a total of 350 potential salivary proteins were identified. Given that salivary proteins usually contain signal peptides and lack transmembrane domains, a total of 87 proteins were predicted to be secreted into soybean during *R. pedestris* feeding (Fig. 1A). These genes, encoding proteins with signal peptides (SPs), were transiently expressed in the leaves of *N. benthamiana.* GFP was used as a negative control and *Phytophtora infestans* INF1 (Wang et al., 2019) as a positive control. As shown in Fig. S1A-D and Fig. 1B, the full length RPH1 protein (*Riptortus pedestris* HAMP 1) induced cell death. Furthermore, when ROS production was measured, the results consistently confirmed that RPH1 triggered cell death and a ROS burst in *N. benthamiana* leaves (Fig. 1C).

**Fig. 1.**
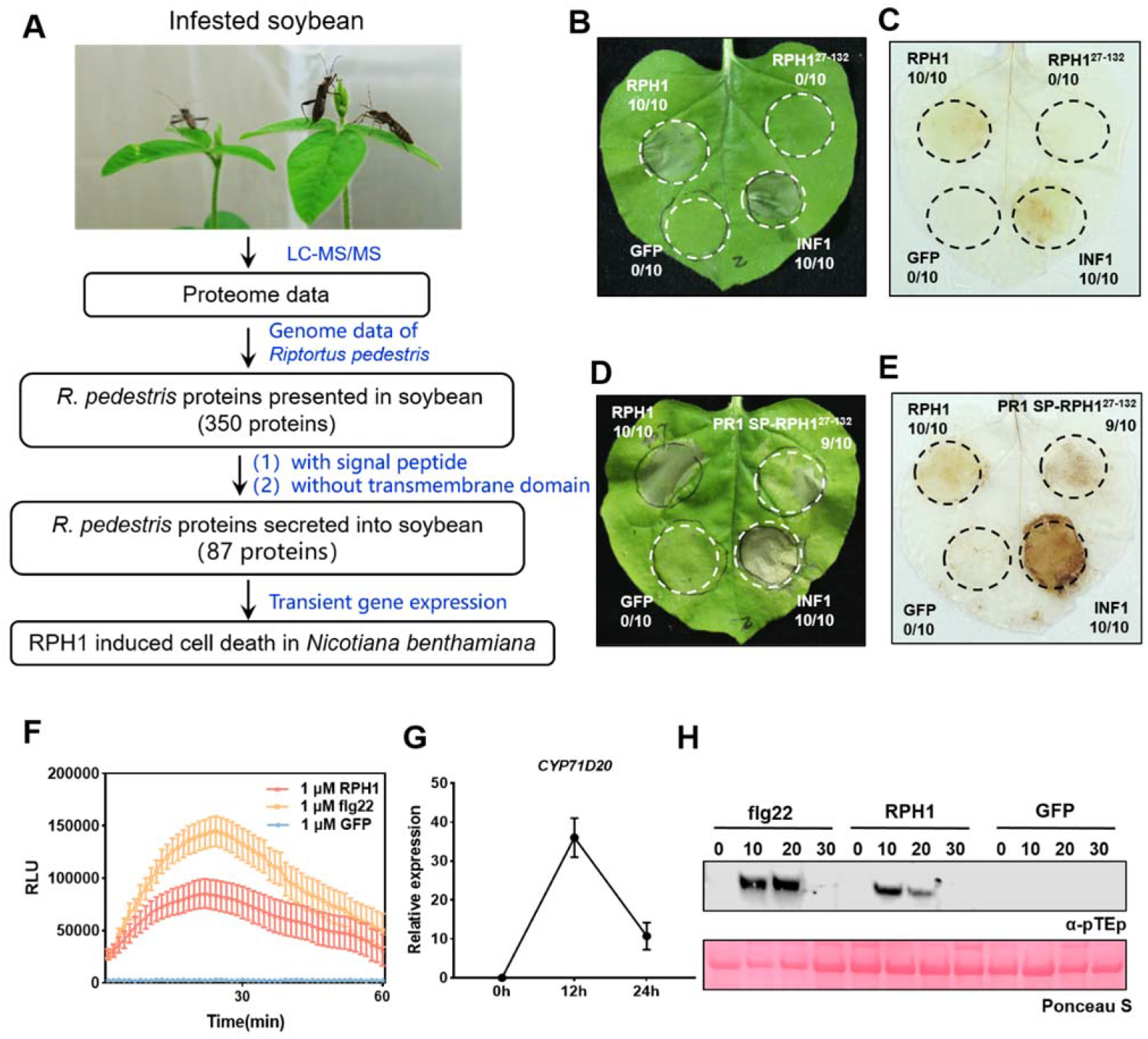
RPH1 from *Riptortus pedestris*, a candidate HAMP, triggered a variety of PTI responses in *N. benthamiana*. **(A) Bioinformatics pipeline for identification of candidate HAMPs. (B and D) Transient expression of RPH1, RPH1^127-132^, and PR1 SP-RPH1^27-132^ in the leaves of *N. benthamiana***. GFP was used as a negative control and INF1 was a positive control. The ratios in the circles represent the number of samples inducing cell deaths to the total number of experiments. Photographs were taken 5 d post-infiltration. **(C and E) 3, 3′-diaminobenzidine (DAB) staining of reactive oxygen species (ROS) accumulation induced by RPH1, RPH1^27-132^, and PR1 SP-RPH1^27-132^ 2 d after infiltration. (F) RPH1 triggered ROS burst.** Flg22 and GFP protein were used as positive and negative controls, respectively. ROS production was measured with a luminol-based assay. Mean RLU (Relative Luminescence Unit) (±SD) are shown (n=14). **(G) RPH1 upregulated the expression of *CYP71D20* gene**. Relative expression was quantified by reverse qRT-PCR using *Nbactin* as a reference gene. Bars indicate mean fold changes ±SD. **(H) RPH1 Induced MAP kinase activity.** *N. benthamiana* plants were treated with 1 μM RPH1 protein at the indicated times, and then MAPK activity was determined by immunoblot with α-pTEp antibody. Protein loading was indicated by Ponceau S staining for Rubisco protein.

RPH1 encodes 132 amino acids with a signal peptide (SP) at the N-terminus, and contains a putative MBF2 transcription activator domain (Fig. S1E). We next deleted the secretion signal and found that RPH1 lacking SP (RPH1^27-132^) failed to trigger cell death and ROS (Fig. 1B and C). By contrast, when the SP of RPH1 was replaced with the pathogenesis-related protein 1 (PR1) signal peptide, cell death and ROS inducing activity were restored (PR1 SP-RPH1^27-132^; Fig. 1D and E). Western blot analysis confirmed that INF1, GFP, RPH1, RPH1^27-132^ and PR1 SP-RPH1^27-132^ were expressed with the predicted sizes in *N. benthamiana* (Fig. S1J).

We expressed His-tagged RPH1 in *Escherichia coli* and purified the resultant recombinant protein (Fig. S2A). Different concentrations of purified recombinant RPH1 were infiltrated into the leaves of *N. benthamiana*. As shown in Fig. S2B, RPH1 protein induced cell death in a concentration-dependent manner. We further conducted a quantitative luminol-based assay for monitoring ROS production on leaf disks of *N. benthamiana*. The 22-amino-acid peptide flg22 from bacteria, a well-known PAMP, was used as a positive control (Yu et al., 2017). As shown in Fig. 1B, the ROS burst triggered by RPH1 peaked after 7 min, the same as with flg22. The amount of ROS production increased in response to higher concentrations of RPH1 (Fig. S2C). In summary, RPH1 stimulates cell death and ROS in *N. benthamiana* in a concentration dependent manner.

Marker genes frequently used for monitoring PTI, namely *CYP71D20*, *WRKY7*, *WRKY8*, *Acre31*, and *PR1,* were examined to assess the elicitor function of RPH1 (Xu et al., 2022). As shown in Fig. 1C and Fig. S2D-G, *CYP71D20*, *WRKY7*, *WRKY8*, *Acre31*, and *PR1* were significantly up-regulated in *N. benthamiana* leaves upon RPH1 treatment (Fig. S2G). We further detected MAPK activation after RPH1 treatment by immunoblotting with the α-pTEpY antibody. As shown in Fig. 1D, RPH1 activated MAPK phosphorylation at 10 and 20 min in *N. benthamiana*. To test the effectiveness of the defense response induced by RPH1, leaves of *N. benthamiana* were pre-treated either with 1 μM RPH1 or with GFP protein, and then 24 h later were inoculated with mycelium of *Phytophthora capsici*. After 48 h of infection (hpi), the leaf area pretreated with RPH1 significantly developed smaller lesions in comparison with GFP treatment (Fig. S2H and I). Measurement of biomass in the inoculated *N. benthamiana* leaves using quantitative genomic DNA PCR (qPCR) revealed that RPH1 reduced *P. capsici* growth significantly compared with the GFP control (Fig. S2J), supporting that RPH1 enhanced plant immunity against pathogens.

### RPH1-induced immune responses require co-receptors BAK1 and SOBIR1

BAK1 is a key co-receptor of PRRs that is indispensable for signaling by both RLKs and RLPs (Chinchilla et al., 2009). To determine whether *BAK1* participates in RPH1 perception or signaling, we generated virus-induced gene silencing (VIGS) constructs targeting *BAK1* expression in *N. benthamiana*, and the silencing efficiency was confirmed using qRT-PCR (Fig. S3A). Our results revealed that RPH1 or INF1-triggered cell death and ROS in TRV-*GUS* lines was impaired in *BAK1*-silenced plants (Fig. 2A-C). *CYP71D20* and *WRKY7* transcripts were significantly reduced in TRV: *BAK1* lines compared with TRV: *GUS* lines (Fig. 2D and E). RPH1-induced MAPK activation was also altered in TRV-*BAK1* lines compared with TRV: *GUS* lines (Fig. 2K). Thus, RPH1-triggered immune responses were dependent on *BAK1*.

**Fig. 2.**
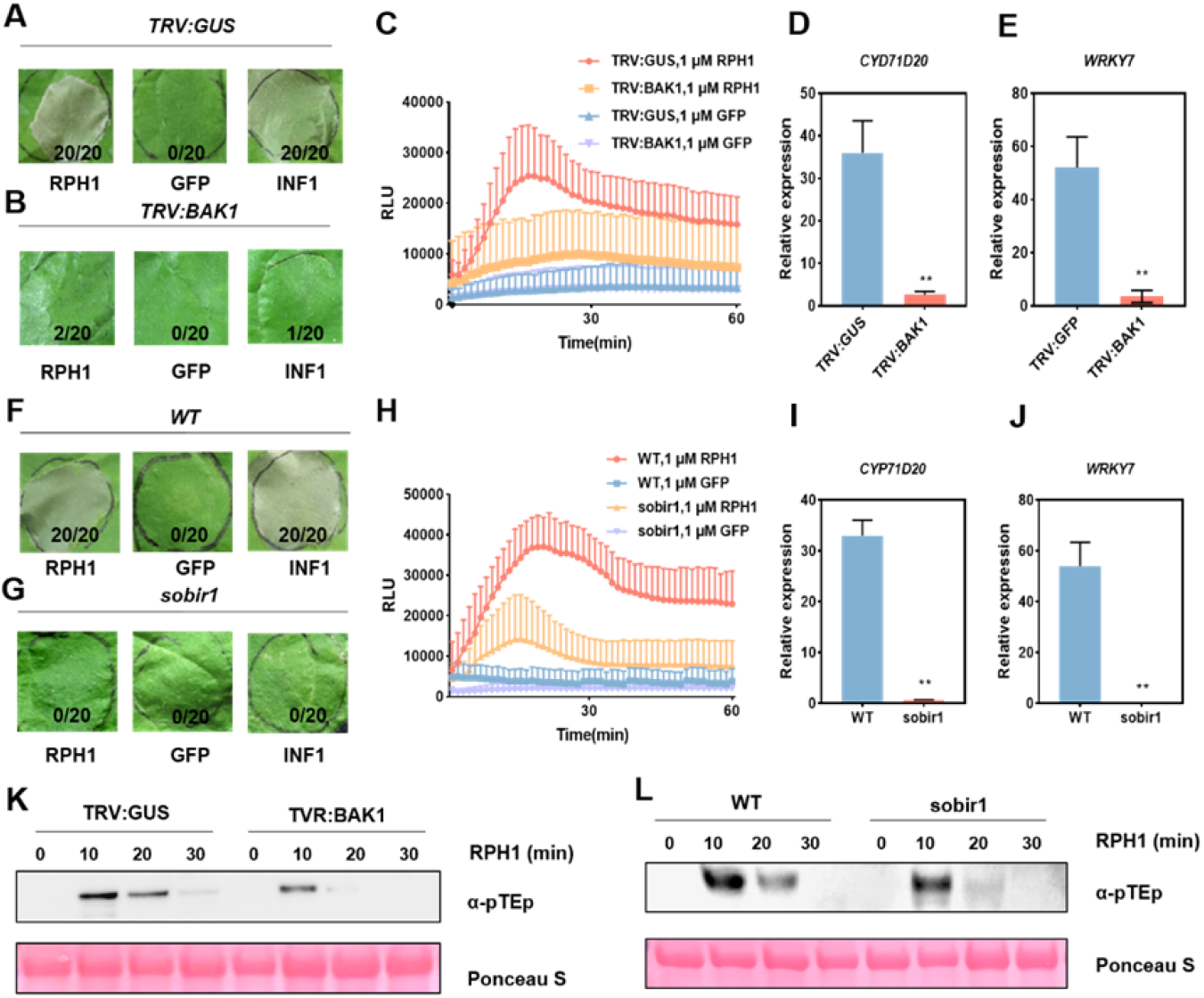
RPH1-induced immune responses require co-receptors BAK1and SOBIR1. **(A) RPH1-induced cell death observed in *TRV:GUS* plants was compromised in *TRV:BAK1*-silenced plants**. GFP and INF1 were used as a negative and positive control, respectively. Photographs were taken 5 h post agroinfiltration. **(B) RPH1 failed to induce ROS bursts in *BAK1*-silenced tissues**. Mean RLU (±SD) are shown (n=14). **(C) The upregulation of *CYP71D20* and *WRKY7* transcripts triggered by RPH1 was lost in *BAK1*-silenced plants**. Error bars indicate the mean fold changes ± SD (**P < 0.01, Student’s t-test). **(D) RPH1 failed to induce cell death in a *sobir1* knockout mutant**. Wild type (WT) *N. benthamiana* was used as a positive control. **(E) RPH1-induced ROS curve in *N. benthamiana* WT was compromised in a *sobir1* mutant. (F) RPH1 failed to upregulate the gene expression of *CYP71D20* or *WRKY7* in a *sobir1* mutant**. Error bars indicate the mean fold changes ± SD (**P < 0.01, Student’s t-test). **(G and H) MAPK activation induced by RPH1 was reduced in *BAK1*-silenced and *sobir1* mutant plants.**

RLPs specifically interact with co-receptor SOBIR1 for triggering downstream signaling (Liebrand et al., 2014). We used a previously generated *sobir1* CRISPR/Cas9 mutant to test whether it is involved in the immune induction of RPH1 (Wang et al., 2022). The results revealed that RPH1 failed to induce cell death, ROS burst, or upregulation of defense genes expression in this mutant (Fig. 2F-J). MAPK activation was also markedly reduced in sobir1-mutant plants compared with WT plants (Fig. 2L). Therefore, immune responses triggered by RPH1 require both PRR-co-receptors, BAK1 and sorbir1, indicating that RPH1 might be recognized by an RLP in the apoplast.

### RPH1 induces PTI responses in soybean

To assess the ability of RPH1 to trigger PTI in soybean leaves, we sprayed 1 μM RPH1 protein on soybean leaves, and then measured ROS production. The results showed that RPH1 protein induced a strong ROS burst in soybean leaves, and the induction by RPH1 was similar to that triggered by flg22 (Fig. 3A). The accumulation of ROS was positively correlated with RPH1 protein concentration (Fig. 3B). In addition, the relative expression of *GmPR1* was increased approximately ten-fold by RPH1 compared to GFP (Fig. 3C). Plant phytohormones including JA, SA and ET have been implicated in early responses against herbivorous insects (Schuman and Baldwin, 2016). We investigated the effects of RPH1 on the biosynthesis of phytohormones. As shown in Fig. 3D-F, spraying of RPH1 protein on the soybean leaves induced a significant increase in the content of JA and ACC, the precursor of ET biosynthesis. On the other hand, the biosynthetic pathway of SA was reduced by RPH1 treatment (Fig. 3F). Overall, RPH1 triggered ROS production, upregulation of *PR1* gene expression, and JA and ET hormone biosynthesis in soybean.

**Fig. 3.**
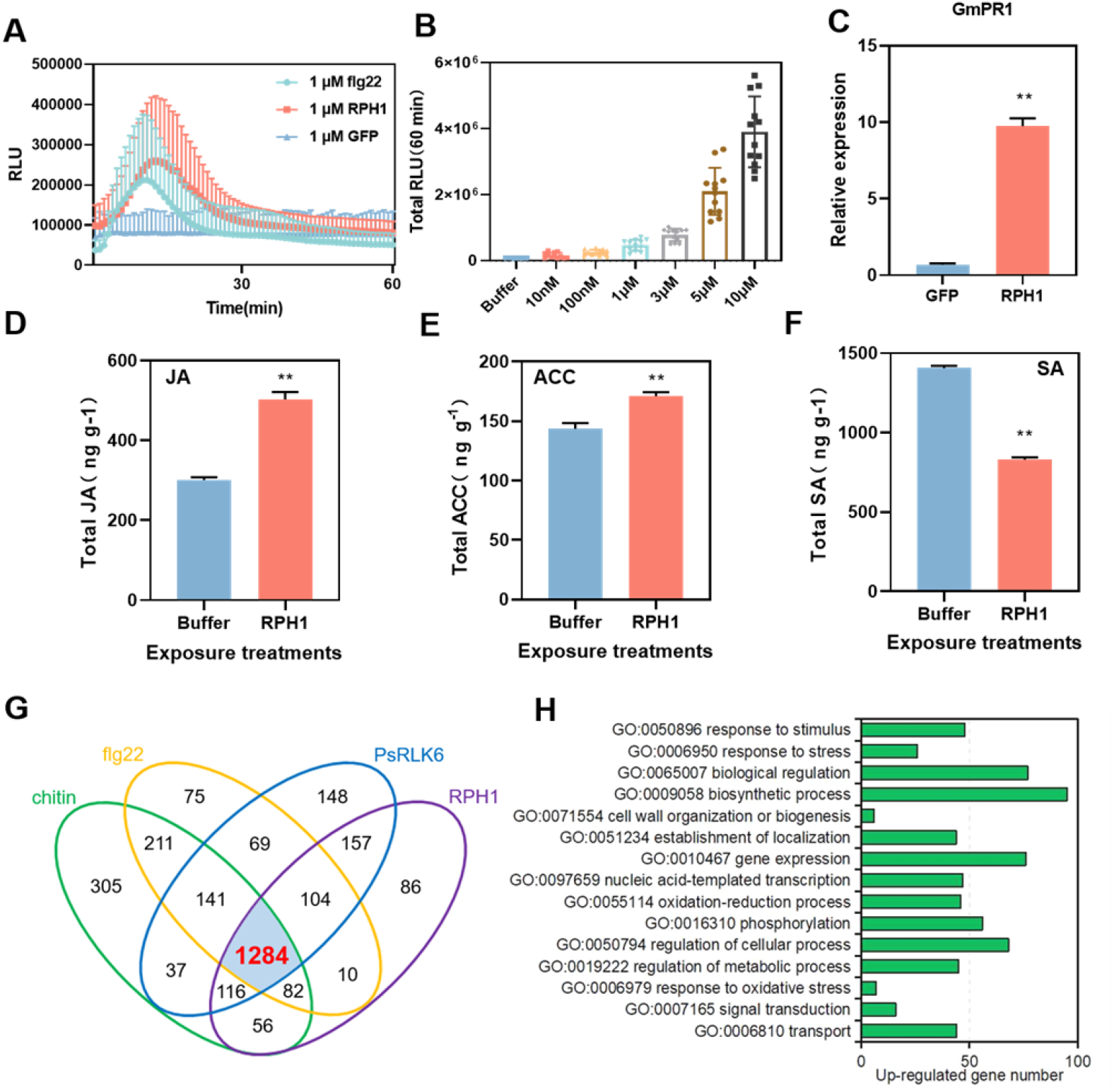
RPH1 induces PTI responses in soybean. **(A) RPH1 induced ROS burst in soybean leaves.** The indicated proteins, including the control GFP, were sprayed onto leaves. ROS production was measured with a luminol-based assay over time. Mean RLU (Relative Luminescence Unit) (±SD) are shown (n=14). **(B) RPH1 triggered ROS production in a concentration-dependent manner.** The indicated concentrations of RPH1 protein, or a buffer control (Bfr), were sprayed onto leaves. The total relative luminescent units were detected over a 60 min period. The results were obtained from two independent experiments with 12 leaf disks in each. **(C) RPH1 induced the upregulation of *GmPR1* transcript.** 1 µM RPH1 protein or the control GFP, were sprayed onto leaves, and RNA was extracted after 12 hours. Relative transcript levels were quantified by RT-qPCR with *GmELF1β* as a reference. Bars indicate mean fold changes ± SD (**P < 0.01, Student’s t-test). **(D-F) RPH1 increased the biosynthesis of JA and ACC while reducing the content of SA.** The contents of JA, ACC and SA were measured after 24 h of RPH1 or buffer treatment. Statistical differences were calculated from three independent experiments (**P < 0.01, Student’s t-test). **(G) A four-set Venn diagram showing numbers of up-regulated genes following PAMP/HAMP treatment.** RNA-Seq data were generated from the total RNAs isolated from soybean leaves which were pretreated with 1 µM Chitin, flg22, PsRLK6 and RPH1 proteins for 12 hours. **(H) GO annotation terms of 1284 up-regulated genes in the biological process groups**.

RNA-Seq assays were further conducted to analyze differentially expressed genes (DEGs) induced by different known PAMPs and RPH1. 1 µM bacterial flg22 (Felix et al., 1999), fungal chitin (a homopolymer of β-1,4-linked GlcNAc from fungal cell walls) (Shinya et al., 2015), *Phytophthora sojae* PsRLK6 (Pei et al., 2023) and RPH1 protein were used to treat soybean leaves for 12h, and then RNAs were extracted for RNA-Seq assays. Overall, 1,284 genes were up-regulated among three PAMPs and RPH-1 (Fig. 3G), and Gene Ontology (GO) enrichment analysis revealed that 1284 up-regulated genes were enriched in biological processes related to plant immunity (Fig. 3H). As shown in Fig. S4A, a massive number of defense-related genes were significantly upregulated by treatment of RPH1 including pathogenesis-related genes, ROS-producing genes, transcription factors, plant hormone biosynthesis and signaling-related genes, such as abscisic acid (ABA), JA and ET pathway genes, secondary metabolism, cell wall pathway genes. Thus, results demonstrated that RPH1, analogous to other known PAMPs, activated a wide variety of PTI responses in soybean.

To examine disease resistance conferred by RPH1, etiolated soybean hypocotyls were treated with RPH1 and control GFP proteins, and then inoculated with mycelium of *P. sojae*, a dominant oomycete pathogen of soybean. Soybean hypocotyls pretreated with RPH1 developed significantly weaker symptoms, much shorter lesion lengths and less *P. sojae* biomass in comparison with GFP control. Thus, our results demonstrated that RPH1 enhanced soybean resistance against *P. sojae* (Fig. S4B-D).

### RPH1 enhances soybean resistance against insect feeding

To assess the impact of RPH1 treatment on insect feeding, we placed 60 *R. pedestris* adults and fourth-nymphs into cages each containing two soybean plants (one pretreated with 1 μM RPH1 protein solution and another one with an equal volume of 1 x PBS buffer). As shown in Fig. 4A, the majority of the *R. pedestris* insects preferred to feed on the soybean plants pretreated with buffer instead of RPH1. An average of 19 insects fed on soybean plants without RPH1 treatment while only 5 insects infested plant with RPH1 protein (Fig. 4B). Statistical analysis indicated that the numbers of insects landing on the soybean plants treated with RPH1 and buffer were significantly different.

**Fig. 4.**
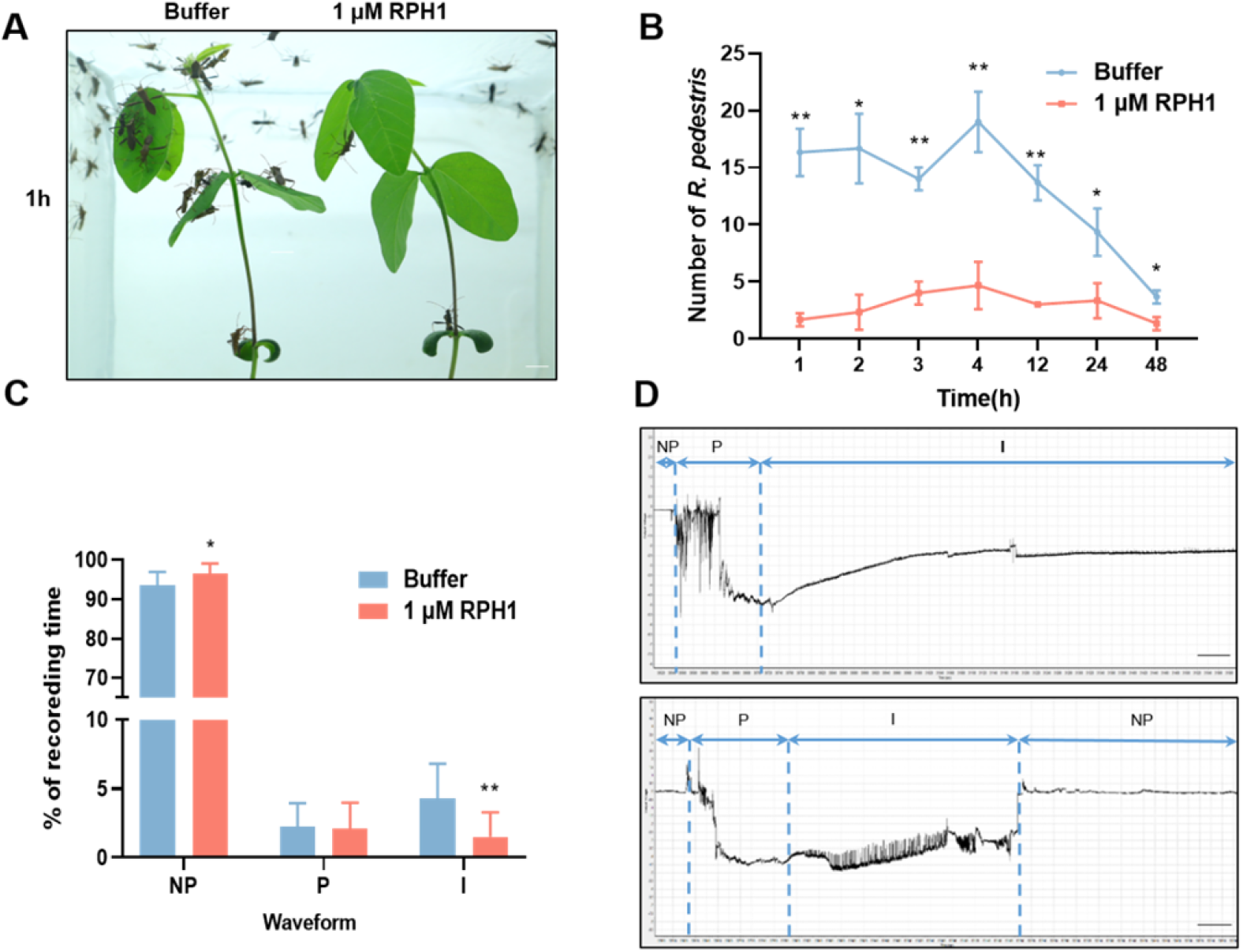
RPH1 enhances soybean resistance against insect feeding. **(A) *R. pedestris* preferred to feed on soybean plants without RPH1 protein treatment**. Photograph was taken at 1 h after placing insects on soybean which were sprayed with indicated proteins. Bar = 1 cm. **(B) Insect numbers of *R. pedestris* feeding on soybean plants were recorded at different times**. Error bars indicate means ± SD (*p < 0.05; **p < 0.01, Student’s t-test). **(C) RPH1 significantly reduced ingestion time of *R. pedestris* on soybean plants**. Insect feeding behaviors were recorded by electrical penetration graph (EPG). NP indicates noneprobing; P represents pathway and I indicates ingestion. Statistical differences were calculated from twenty replicates (*p < 0.05; **p < 0.01, Student’s t-test). **(D) Waveforms for *R. pedestris* feeding on soybean leaves with and without RPH1 treatments were recorded using EPG monitor during 1 h**. The upper figure was generated when *R. pedestris* was feeding on soybean leaves treated with 1 x PBS buffer; the lower figure was recorded during insect feeding on soybean with 1 μM RPH1 protein treatment. Vertical numbers indicate output voltage; horizontal numbers indicate the recording time (second).

We further monitored the feeding behaviors of *R. pedestris* on soybean plants using electropenetrography (EPG). Waveforms of NP (noneprobing), P (pathway), and I (ingestion) were characterized as we previous published (Jin et al., 2022). The results revealed that on RPH1 pretreated leaves, *R. pedestris* spent longer times standing still or walking than on buffer treated leaves. Furthermore, the ingestion period was significantly shorter on RPH1 pretreated soybean leaves than that on buffer-treated leaves (Fig. 4C and D). Statistical analysis demonstrated that in a total of 16 recording hours, the ingestion time on RPH1-treated *G. max* leaves was 147 min, while it was 411 min on buffer-sprayed leaves (Fig. S4E). Collectively, our results revealed that RPH1 treatment strengthens plant deterrence of *R. pedestris* feeding.

### RPH1 is a conserved HAMP in herbivorous insects

MAMPs are molecular signatures typically conserved in whole classes of microbes (Boller and Felix, 2009). We analyzed the distribution of RPH1 homologs in various organisms by BlastP (E-value < 1e-5). RPH1-like sequences were only present in *Streptomyces clavuligerus* among bacteria but were widely distributed within the class of Insecta, indicating that RPH1 proteins are common across insect taxa. Using genome data downloaded from http://www.insect-genome.com/, RPH1 homologs were identified from 25 species which all contained signal peptides. As shown in Fig. 5, RPH1 homologs are distributed in Hemiptera, Lepidoptera, Diptera and Thysanoptera within Insecta. Importantly, the majority of herbivorous insects belong to these four orders (Bonaventure G, 2012). Thus, the results revealed that RPH1 and its homologs play important role in eliciting plant defense responses to herbivorous insects.

**Fig. 5.**
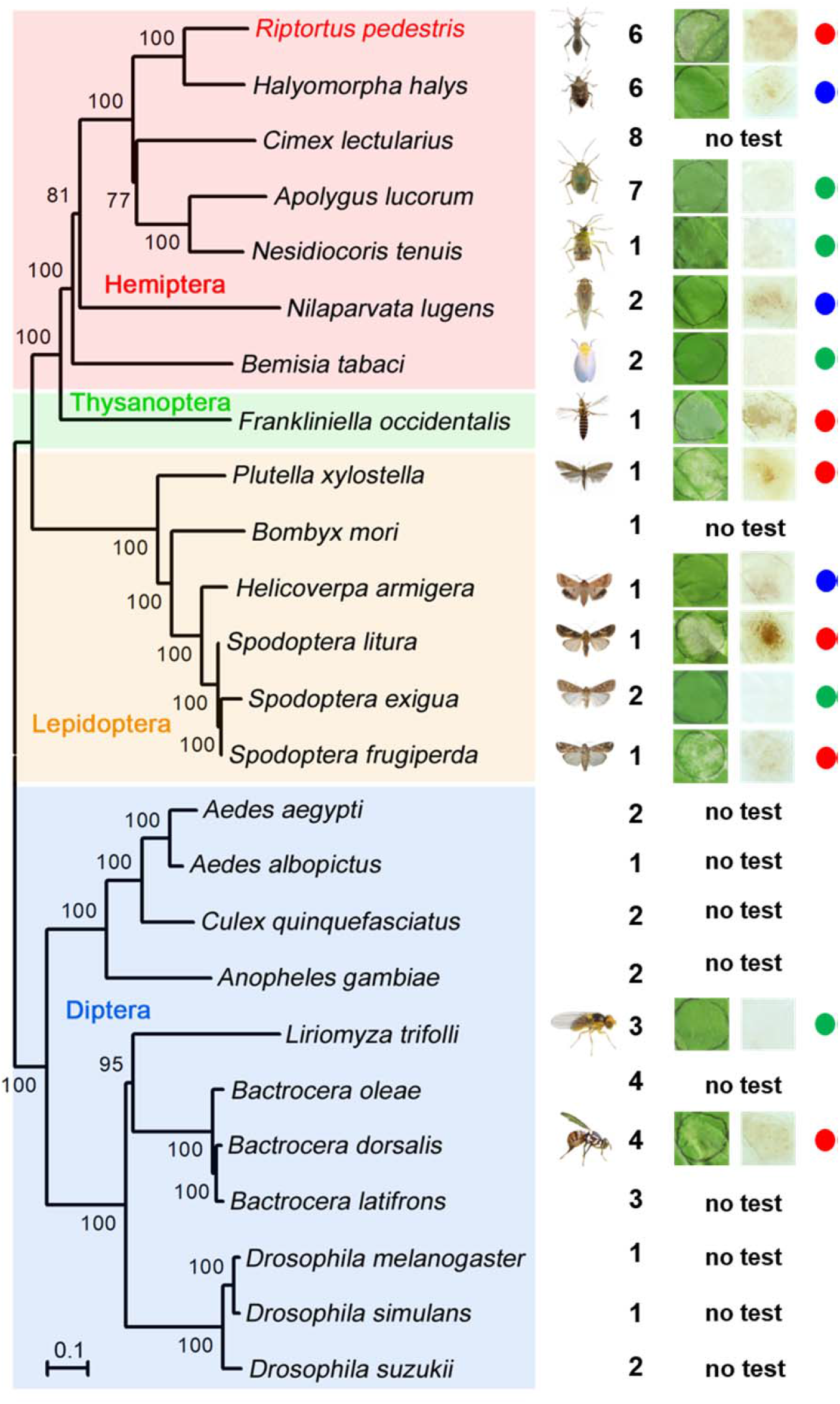
RPH1 is a conserved HAMP in herbivorous insects. RPH1 was distributed in Hemiptera, Lepidoptera, Diptera and Thysanoptera within Insecta. The multigene phylogenetic tree was conducted by MEGA 11 according to the concatenated sequences of single-copy core proteins derived from selected insects. The number represents the number of RPH1 homolog gene. Red dot indicates these proteins induced cell death and ROS burst; blue dot represents that these proteins only induced ROS accumulation while green dot represents that these proteins did not induce cell death or ROS accumulation in *N. benthamiana.* “No test” means that these genes were not investigated.

### *R. pedestris* has evolved paralogous genes to evade plant immunity triggered by **RPH1**

We finally explored whether *R. pedestris* have evolved paralogs to evade PTI triggered by RPH1. A total of 5 paralogs (named RPH1L1, 2, 3, 4, and 5) were obtained and all contains SPs (Fig. S6A). A phylogenetic tree was constructed using RPH-1 and the five paralogous gene sequences (Fig. 6A). Similar to RPH1, RPH1L1 with signal peptides also induced cell death, ROS burst, upregulation of defense marker genes *WRKY7*, *CYP71D20*, *Acre31* and *PTI5*, and enhanced plant resistance against *P. capsici* in *N. benthamiana* (Fig 6A and Fig. S6B-H). Thus, RPH1L1 is also a HAMP in analog to RPH1.

**Fig. 6.**
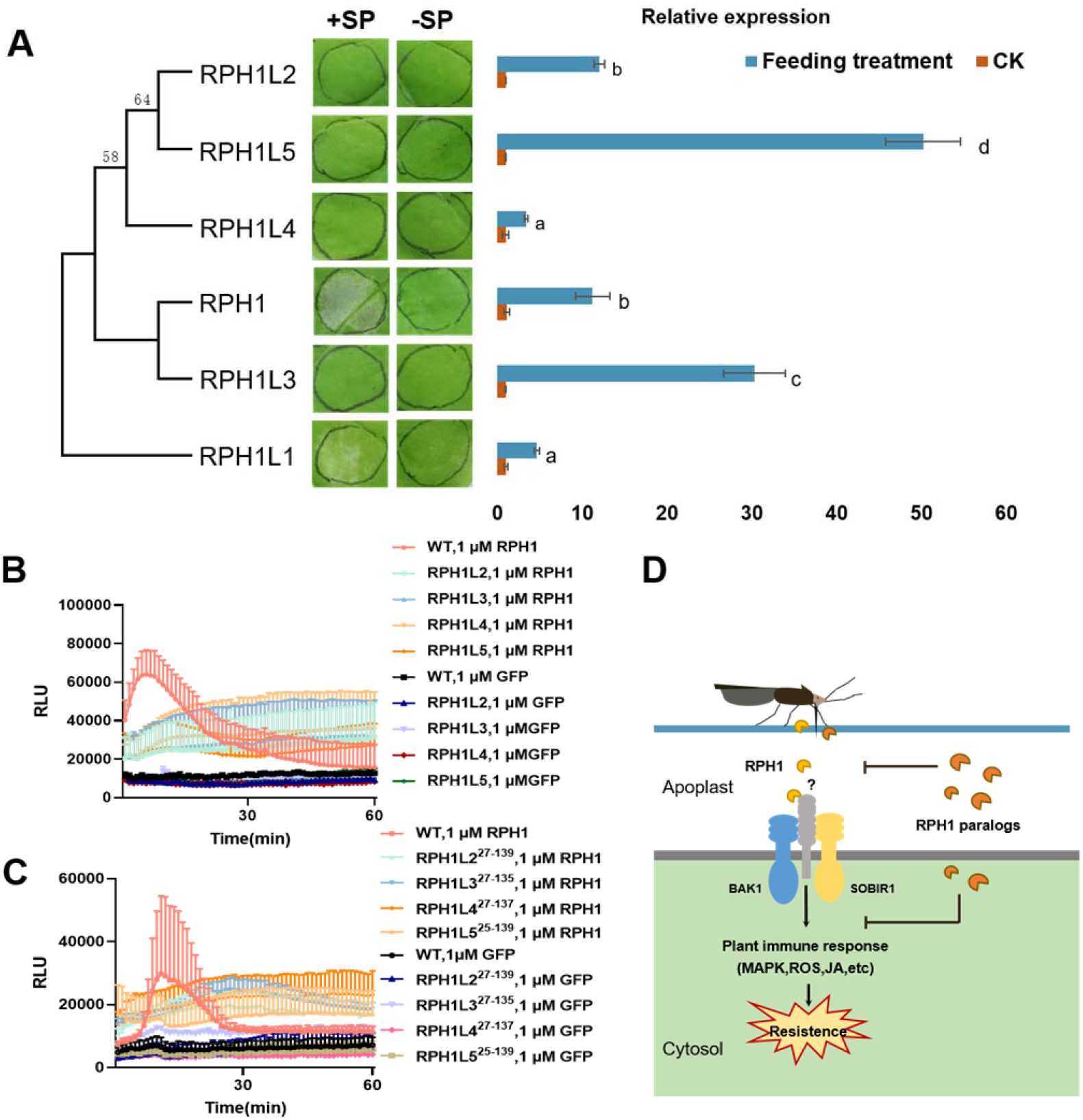
*R. pedestris* evolved paralog genes to evade plant immunity triggered by RPH1. **(A) Phylogenetic tree generated using RPH1 and its paralogous proteins, *N. benthamiana* expression phenotypes of the six genes and relative transcript levels in the insect with and without soybean feeding**. SP: signal peptide. A total of six adults *R. pedestris* insects were divided into two groups. One group was fed on soybean while the other was starved for 24 h. Then expression levels of RPH1 and five paralog genes were measured by qRT-PCR using RNAs from insect whole bodies as templates. Significant differences are calculated from three replicates and indicated by different letters (P < 0.01; Duncan’s multiple range test). **(B) Expression of four paralogs with SPs suppressed the ROS burst triggered by exogenous RPH1**. Mean RLU (Relative Luminescence Unit) (±SD) are shown (n=14). **(C) Expression of four RPH1 paralogs without SPs suppressed the ROS burst triggered by exogenous RPH1.** The results were obtained from two independent experiments. Mean RLU (Relative Luminescence Unit) (±SD) are shown (n=14). **(D) A model illustrating how *R. pedestris* may have evolved paralogs to evade plant immunity triggered by RPH1.** Arrows indicate positive regulation. Barred lines represent negative regulation. Question mark represents an undefined mechanism.

While the rest four paralogs (RPH1L2, RPH1L3, RPH1L4 and RPH1L5) failed to elicit cell death with or without signal peptides (Fig. 6A). We expressed these four paralog genes in *N. benthamiana* for 24 hours, and then infiltrated the full length of RPH1. As shown in Fig 6B and Fig. S6I-J, the cell death and ROS burst induced by RPH1 were disrupted by the expression of four paralog proteins in the apoplast. The upregulation of defense gene WRKY7 was suppressed in a similar manner (Fig. S6K). All proteins were expressed with the expected sizes (Fig. S6O). To further confirm that four paralogs interrupted the RPH1 perception in the extracellular space, we test the effect of paralogs on the ROS triggered by flg22. Our result revealed that four paralogs with SPs failed to suppress the ROS accumulation induced by flg22 (Fig. S7A). Hence, our results suggested that four paralogs of RPH1 might disrupt the perception of RPH1 in the extracellular space, resulting in the inactivation of plant PTI responses.

As shown in Fig. 6C, four paralogs without SPs inhibit the cell death and ROS curve triggered by RPH1. RPH1-induced upregulation of WKRY7 was also subverted by four paralog proteins (Fig. S6L-N). Fig. S6P shown that all proteins were successfully expressed. The ROS curve induced by flg22 was also inhibited by four paralogs (Fig. S7B). Thus, our results revealed that RPH1L2, RPH1L3, RPH1L4 and RPH1L5 inhibit the PTI responses induced by RPH1 and flg22 inside plant cells. Finally, we investigated the expression profiles of RPH1 and paralogous genes in *R. pedestris* while feeding on soybean leaves. As shown in Fig. 6A, the relative expression levels of RPH1L2, RPH1L3 and RPH1L5 were significantly increased in comparison of RPH1 and RPH1L1 after insect feeding. Especially, RPH1L3 and RPH1L5 transcripts were increased multiple folds than RPH1 and RPH1L1.

## Discussion

M15ore than sixty PAMPs have been identified from bacteria, fungi and oomycetes (Boutrot and Zipfel, 2017). Compared to the large number of PAMPs from plant pathogens, the research on HAMPs has lagged behind. Several HAMPs including vilicitin, inceptins, GOX and lipases have been isolated from chewing insects by purification of secreted salivary proteins (Acevedo et al., 2015). A few salivary proteins from piercing-sucking insects have been identified to activate plant defenses (Snoeck et al., 2022), whereas none of them was recognized by plant PRR in the apoplast. In this study, the large size of the insects and heavy feeding on plants enabled us to identify *R. pedestris* proteins secreted into soybean tissue using LC-MS/MS. Taking advantage of the large scale elicitor screening platform provided by *N. benthamiana*, one *R. pedestris* protein RPH1 was identified to induced cell death in the extracellular space. Further results confirmed that RPH1 activated a repertoire of PTI responses in non-host plant *N. benthamiana* and host plant soybean including ROS, MAPK activation, upregulation of defense genes and accumulation of hormones such as JA and ET. Immune defenses triggered by RPH1 required PRR co-receptors *BAK1* and *SOBIR1*, indicating that RPH1 might be recognized by a plant RLP.

Herbivorous insects mainly belong to the orders Lepidoptera, Coleoptera, Hemiptera, Thysanoptera, and Diptera (Bonaventure G, 2012). Only chewing insects such as grasshoppers within Orthoptera are classified into Polyneoptera, while the rest fall into Eumetabola. PAMPs are often highly conserved molecules with signatures characteristic of a whole class of microbes (Boutrot and Zipfel, 2017). In this study, RPH1 identified from the mirid bug *R. pedestris* is highly conserved within Eumetabola, and has homologous genes in species within the orders Diptera, Hemiptera, Thysanoptera, and Diptera. The majority of selected species from the four orders activated the first-layer immune response, PTI, in the model plant *N. benthamiana*, revealing their critical role in plant pattern signal recognition. Homologs of RPH1 are not only present in herbivorous pests, but also are widely distributed in blood-sucking mosquitoes, such as *Anopheles gambiae*, *Aedes albopictus* and *Culex quinquefasciatus*. Owing to the striking similarity of plant and animal innate immunity (Nurnberger et al., 2004), RPH1 homologs in mosquitoes might also serve as molecular patterns recognized by human cells. Our results also demonstrated that homologs of RPH1 from both chewing and piercing-sucking insects trigger plant immunity. Characterized HAMPs including Volicitin and inceptins are only conserved in chewing insects (Schmelz et al., 2006; Alborn et al., 2007). Our study provides further evidence that herbivorous insects share common pattern molecules despite their different mouths and feeding behaviors.

We found that host-adapted *R. pedestris* have evolved four paralogs of RPH1 (RPH1L2, 3, 4, 5) to evade plant PTI responses triggered by HAMP RPH1. Within plant cells, the four paralogs inhibit downstream signaling of PTI activated by RPH1 to evade plant immunity, presumably via the MBF2 transcription activator domain that occupies most of the length of RPH1 and its paralogs. In the apoplastic space, four paralogs might disrupt the perception of RPH1 to disarm PTI activation. This novel adaptation strategy is similar to the decoy model of PsXEG1, which was produced by the soybean pathogen *phytophthora sojae*. Soybean produced a glucanase inhibitor protein, GmGIP1 that binds to PsXEG1, an essential virulence effector of *P. sojae*, to block its contribution to virulence. To counter defense, *P. sojae* secretes a paralogous PsXEG1-like protein, PsXLP1 that binds to GmGIP1 more tightly than does PsXEG1, to freeing PsXEG1 (Ma et al., 2017). After *R. pedestris* infestation, HAMP RPH1 and RPH1L1 upregulation was significantly less than that RPH1L2, 3, 5 (Fig. 6D); possibly to reduce their recognition by the plant immune system.

## Materials and methods

### Insects, plants and microbial materials

A laboratory strain of *R. pedestris* from a previous article was used for this study (5). Nymphs and adults were fed with potted soybean plants and dried seeds. *N. benthamiana* and soybean strains (cv. Williams and Hefeng47) were cultivated at 25 °C and 60% relative humidity under a 16:8h (light:dark) photoperiod. The *Phytophthora capsici* strain LT263 and the *Phytophthora sojae* strain P6497 were cultured at 25°C in the dark on 10% (v/v) V8 juice medium.

### Identification of *R. pedestris* salivary proteins and *Agrobacterium tumefaciens* infiltration

50 *R. pedestris* nymphs and adults were starved for 2 h, and then placed onto soybean seedlings for infestation. After feeding by *R. pedestris* for 48 h, leaves were detached and instantly placed into liquid nitrogen for shotgun liquid chromatograph mass spectrometry (LC-MS/MS). The resulting spectra were searched against a *R. pedestris* proteome database (Huang et al., 2021). The signal peptide of each protein was predicted using SignalP v3.0, and transmembrane helices in proteins were predicted using the TMHMM Server v2.0. Potential protein domains were predicted using the Pfam database (EMBL-EBI).

Candidate effectors were amplified by PCR using *R. pedestris* genomic DNAs or cDNAs as templates. For overexpression in *N. benthamiana*, the candidate effectors were cloned into plant expression vector pBinGFP2 carrying green fluorescent protein (GFP) tag or pBin3HA with HA tag using a ClonExpress II One Step Cloning Kit (Vazyme, NanJing, China). The positive constructs were transferred into *A. tumefaciens* strain GV3101 by electroporation. Recombinant strains of *A. tumefaciens* were cultured in Luria-Bertani (LB) medium at 28°C with shaking at 200 rpm for 36 h. The cells were then washed and re-suspended in infiltration buffer (10 mM MgCl_2_, 500 mM MES, 150 μM acetosyringone) to an optical density (OD) of 0.4-0.6 at 600 nm. The suspensions were infiltrated into *N. benthamiana* leaves using a syringe.

### Reactive oxygen species (ROS) assays

*N. benthamiana* leaves with expressing effectors were immersed in 1 mg/ml DAB (3, 3′-diaminobenzidine) solution (Sigma-Aldrich, USA) at 25°C for 8 h, and then decolorized and preserved in 30% glycerol. Photographs of DAB-stained *N. benthamiana* leaves were taken under natural light.

Leaf disks collected from 4-week-old *N. benthamiana* or soybean leaves were harvested, and then soaked in water to keep in the dark at room temperature. The leaf disks were stained in test buffer containing luminol (123072; Sigma-Aldrich), L012 (120-04891; Fujifilm WAKO, Chuo-Ku, Japan), horseradish peroxidase (P6782; Sigma-Aldrich), and 1 μM flg22 (RP19986; GenScript, Nanjing, China) or 1 µM RPH1 or 1 µM GFP proteins. Luminescence detection was conducted to measure ROS production (BioTek, Beijing, China).

### Western blots

*N. benthamiana* leaves with expression of candidate effectors for 36-48 h were collected to extract total protein (Si et al., 2021). After separation of isolated proteins on 10% SDS-PAGE gels, proteins were transferred to a membrane for immunoblotting using either mouse anti-GFP monoclonal antibodies (1:5,000; Abmart, Shanghai, China) or anti-HA monoclonal antibodies (1:5,000; Abmart, Shanghai, China).

### Expression and purification of RPH1 protein in *Escherichia coli*

GFP or RPH1 with His tag was cloned into the pET-32a vector, and expressed in Rosetta (DE3) *Escherichia coli* cells. After cells were grown in LB medium with 0.1 mM isopropyl β-D-1-thiogalactopyranoside (IPTG) for 12-16h at 18°C, the cell culture was lysed and collected. The recombinant GFP and RPH1 proteins were enriched and purified by affinity chromatography using Ni-NTA Superflow resin (Thermo Scientific). Protein concentrations were calculated based on a standard curve created using bovine serum albumin (BSA) as a substrate.

### RNA extraction and qRT-PCR analysis

*R. pedestris* cDNA was synthesized from total RNA using the HiScript II QRT SuperMix, and qRT-PCR was performed using SYBR Green Master Mix (Vazyme, Nanjing, China). qRT-PCR was performed on an ABI PRISM 7500 real-time PCR system (Applied Biosystems) to obtain melt curves. The relative expression levels of genes were normalized using β-actin as the reference gene. At least three biologically independent replicates were carried out for each sample. Relative expression was calculated using the 2^−ΔΔCT^ method (Livak and Schmittgen, 2001).

### MAPK phosphorylation

Total proteins were extracted from four-week-old *N. benthamiana* leaves after treatment with 1 μM flg22, RPH1 or GFP protein. Isolated proteins were separated on a 12% SDS-PAGE gel for immunoblotting. Phosphorylation of MAPK proteins including MPK3, MPK4, and MPK6 was determined using anti-phospho-p44/42 MAPK antibody (1:5,000, Cell Signaling Technology, Danvers, Massachusetts, USA; #9101) and HRP-conjugated anti-rabbit IgG (1:10,000, Sigma-Aldrich; A6154) as a secondary antibody for immunoblot detection.

### Pathogen infection and insect feeding assays

5 mm diameter *P. capsici* agar plugs were placed onto *N. benthamiana* leaves expressing effectors or control proteins (GFP) for incubation in dark at 25°C. The infected leaves were photographed under UV light and lesion sizes were measured using ImageJ software at 48 hours post-inoculation (hpi). Biomass of *P. capsici* in infected *N. benthamiana* leaves was quantified using genomic DNA qPCR (quantitative real-time PCR) based on a previously described method (Liang et al., 2021).

Soybean etiolated seedlings were obtained by germinating seeds of a susceptible soybean strain (Hefeng 47) in the dark at 25°C for 4 days. 0.5 cm diameter *P. sojae* myceliual plugs were inoculated onto the hypocotyls of etiolated soybean seedlings which had been immersed in 2 mL of 1 µM RPH1 or GFP protein solution for 6 h. Disease symptoms were recorded at 48 hpi and the lesion lengths (cm) were measured. Relative biomass of *P. sojae* were determined by genomic DNA qPCR. At least three replications were performed for each treatment and significant differences were calculated using Student’s *t* test.

For insect preference feeding assay, 60 *R. pedestris* adults and fourth-stage nymphs were starved for 12-16 hours, and then placed in a cage containing soybean plants sprayed with 2 mL of 1 µM RPH1 protein solution or 1x Phosphate Buffered Saline (PBS) buffer (Vazyme, G101). Feeding preference was recorded with a camera continuously from 1 h to 48 h. This experiment was replicated at least three times and significant differences were analyzed using Student’s *t* test.

### Analysis of hormone contents in soybean leaves

After *G. max* leaves were sprayed with 2 mL of 1 µM RPH1 protein solution or 1x PBS buffer (Vazyme, G101) for 12 h, the treated leaf was detached and instantly frozen in liquid nitrogen. Samples were prepared according to the manufacturer’s instructions (Nanjing Convinced-test Technology Co., Ltd, Nanjing, China). 2 μl of sample solution was injected into the reverse-phase C18 Gemini HPLC column. HPLC–ESI–MS/MS analysis was performed by Nanjing Convinced-test Technology Co to determine the contents of JA, SA and ET hormones.

### Insect feeding behavior analysis using EPG

The feeding behavior of the *R. pedestris* on soybean leaves was monitored using the Giga-8 DC-electrical penetration graph (EPG) System. Waveforms were recorded as previously described (Jin et al., 2022). All experiments were repeated at least three times. Twenty insects were consecutively monitored for 16 h for each treatment. Waveform was analyzed using the STYLET+A software (Dong et al., 2023).

### RNA-Seq analysis

Soybean leaves were sprayed with 2mL 1 μM RPH1 protein solution or 1x PBS buffer (Vazyme, G101) for 12 h, and then frozen in liquid nitrogen. Wild-type soybean leaves were used as the control. The total RNAs were extracted using an RNA-simple Total RNA Kit (Zoman Biotech). RNA libraries were constructed and subjected to sequencing using the Illumina HiSeq X Ten platform. The resulting high-quality reads were mapped to the soybean genome using HISAT2 (Kim et al., 2015). To calculate the expression of each soybean gene, the Stringtie tool was used, normalizing it to the fragment per kilobase of exon per million mapped reads (FPKM) value (Pertea et al., 2015). Differentially expressed genes were identified using DESeq2 with criteria of P value < 0.05 and fold change > 2 or fold change < −2 (Love et al., 2014). The MapMan ontology tool was used to obtain an overview of the metabolic pathways in which *G. max* genes were involved (Thimm et al., 2004). Three sets of biological replicates were prepared for each sample.

### VIGS assay in *N. benthamiana*

pTRV1 and pTRV2: BAK1 plasmid constructs were transformed into *A. tumefaciens* GV3101 using electroporation. pTRV2: GUS was used as a negative control while pTRV2: PDS (phytoene desaturase gene) was used as a positive control. The cultured *Agrobacterium* cells harboring pTRV2: BAK1and pTRV1vector were harvested and diluted in the infiltration buffer to an OD_600_ of 0.5, respectively, and then were mixed together in a 1:1 ratio. The cell mixture was then infiltrated into the lower leaves of *N. benthamiana* during four-leaf stage. At about 3 weeks post infiltration of pTRV strain, the third new leaf was used to extract total RNA and conduct qRT-PCR. The NbEF1a was used as an endogenous control. The experiments were repeated at least three times. *A. tumefaciens* strain harboring RPH1 was then infiltrated into the third new leaves of BAK1-slienced plants.

### Bioinformatics analysis of RPH1 homologs and paralogs

To determine the phylogenetic relationships of selected insects, we used BlastP results (E-value < 1e-5) of all protein sequences derived from studied insects were obtained from http://www.insect-genome.com/ and used as input into TribeMCL for clustering. Protein sequence clusters with only one member from each insect were defined as single-copy core proteins. The multigene phylogenetic tree was conducted by MEGA 11 according to the concatenated sequences of single-copy core proteins. To search for RPH1 orthologs in other insects, we used the protein sequence of RPH1 as the query to perform BlastP analysis (E-value < 1e-5) against other insect proteomes. The phylogenetic tree of RPH1 and its homologs was constructed using the neighbor-joining algorithm with 1,000 bootstrap replicates by MEGA 11 software.

## Author contributions

DD, YZ and AX conceived and designed research. JZ, YY, QZ, YM, WL and YZ conducted experiments. YW, YP and GA helped designing research. JZ, GA, and DS analyzed data. AX and JZ wrote manuscript. DD and DY revised manuscript.

## Acknowledgements

We thank Dr. Fajun Chen, Shuwen Wu, Zewen Liu, Ming Zhu and Xiangdong Liu for providing insect materials. We also thank Dr. Jinding Liu for bioinformatics analyses.

## Supplementary Figures

**Fig. S1.**
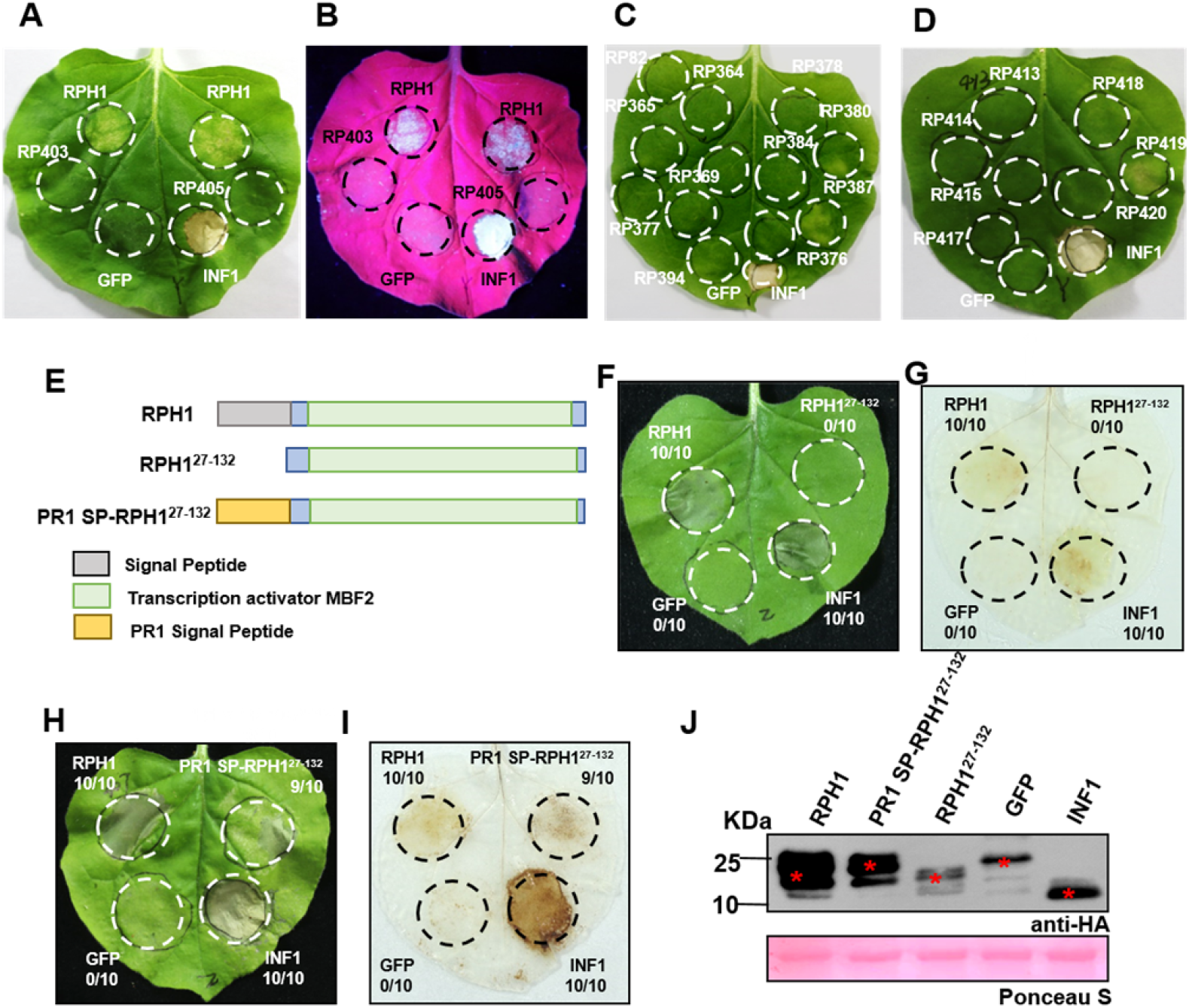
*Riptortus pedestris* RPH1 is an apoplastic elicitor of cell death. **(A-D) Partial screening results of potential *R. pedestris* HAMPs in *N. benthamiana* by agroinfiltration**. Photo B was taken under UV light. **(E) RPH1 gene analysis and construction of PR1 SP-RPH1^27-132^. (F and H) Transient expression of RPH1, RPH1^127-132^, and PR1 SP-RPH1^27-132^ in the leaves of *N. benthamiana***. GFP was used as a negative control and INF1 was a positive control. The ratios in the circles represent the number of samples inducing cell deaths to the total number of experiments. Photographs were taken 5 d post-infiltration. **(G and I) 3, 3 ′-diaminobenzidine (DAB) staining of reactive oxygen species (ROS) accumulation induced by RPH1, RPH1^27-132^, and PR1 SP-RPH1^27-132^ 2 d after infiltration. (J) Western blot detection of expressed proteins using anti-HA antibodies.** HA, hemagglutinin. The red asterisks (*) indicate the predicted protein sizes. Ponceau staining of Rubisco protein (PS) indicated that equal amount of each sample was loaded.

**Fig. S2.**
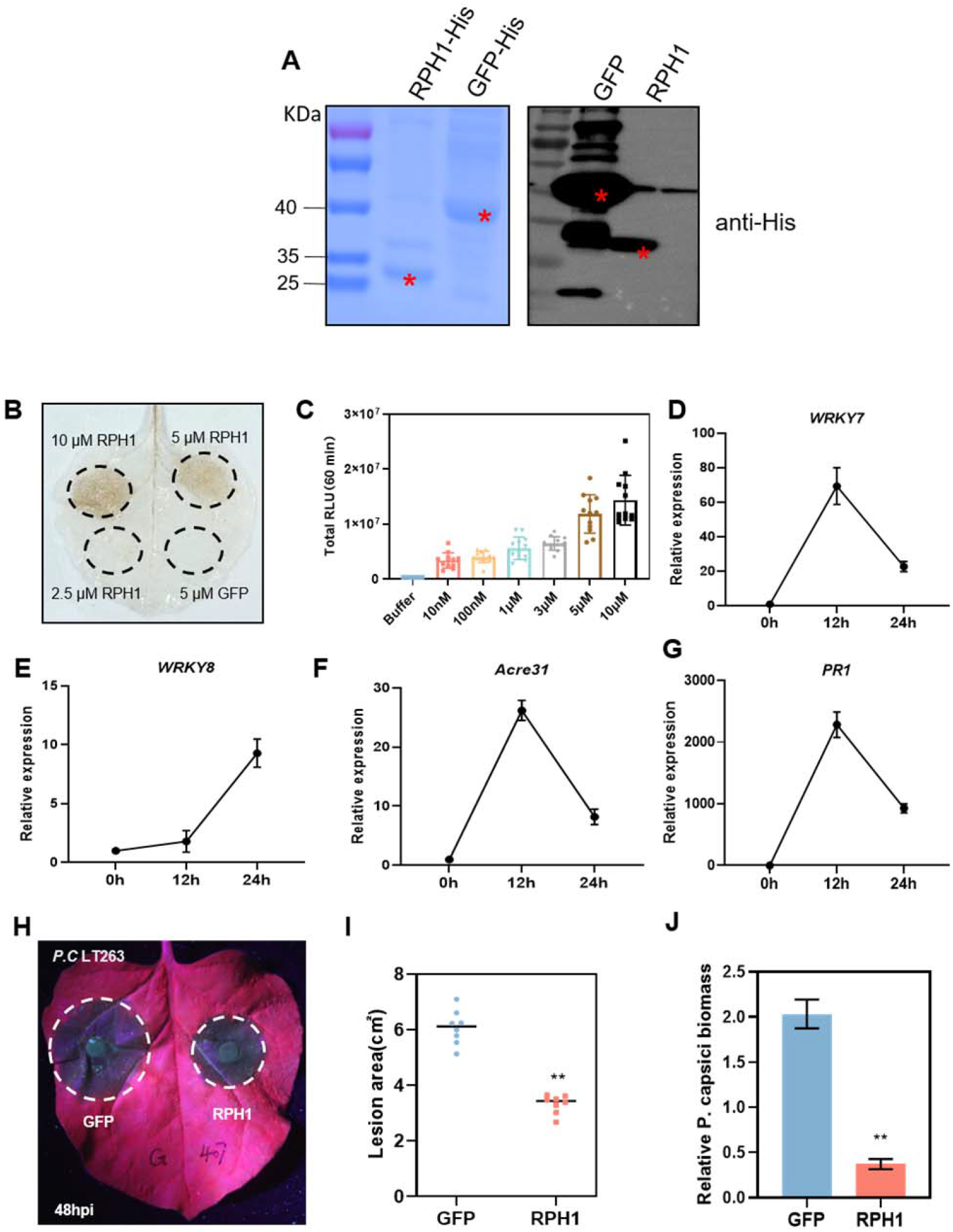
RPH1 triggered a variety of PTI responses in *N. benthamiana*. **(A) Expression of recombinant RPH1 proteins in *Escherichia coli***. GFP was a control. Left figure was a purified RPH1 and GFP proteins gel image. Right figure was western blot detection of RPH1 and GFP proteins with anti-His antibody. The red asterisks (*) indicate the predicted protein sizes. **(B) RPH1 protein induce ROS accumulation in *N. benthamiana* leaves in a concentration-dependent manner**. GFP serves as a control. **(C) RPH1 induced ROS burst in a concentration-dependent manner in *N. benthamiana* leaves**. The relative luminescent unit was continuously measured for 60 mininutes. The results were obtained from two independent experiments, and twelve leaf disks per treatment were used for each experiment. **(D-G) RPH1 triggered the upregulation of PTI marker genes.** Relative transcript levels were quantified by reverse transcription-quantitative PCR (qRT-PCR) using *Nbactin* as a reference gene. Bars indicate mean fold changes ± SD. **(H) RPH1 treatment enhanced plant resistance against *Phytophthora capsici***. Photos of disease symptoms were taken 48 h post inoculation (hpi) under UV light. **(I and J) RPH1 treatment reduced *P. capsici* infection**. For each treatment, lesion areas and relative pathogen biomass were calculated from eight biological replicates. Total DNA of leaves infected by *P. capsici* for 48 h was extracted for quantitative PCR analysis. *NbActin* was used as a reference gene. Error bars indicate means ± SD (**P < 0.01, Student’s t-test).

**Fig. S3.**
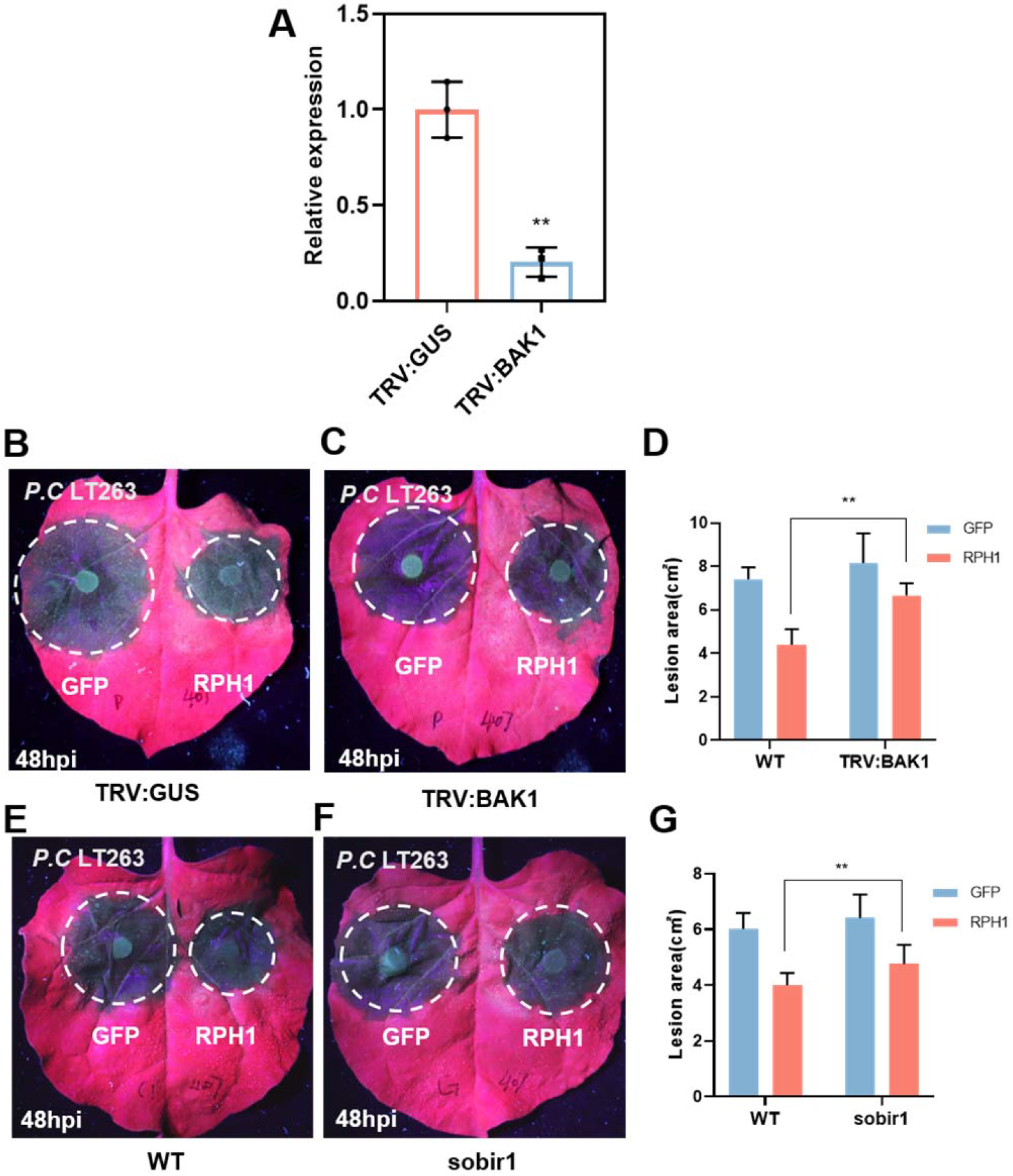
RPH1-induced immune responses require co-receptor BAK1and SOBIR1. **(A) Silencing efficiency of *BAK1* was verified by qRT-PCR.** Error bars indicate the mean fold changes ±SD (**P < 0.01, Student’s t-test). **(B-D) RPH1-induced plant resistance against *Phytophthora capsici* in TRV-GUS lines was compromised in the BAK1-silenced plants**. Photographs were taken 48 hpi under UV light. The lesion area was measured at 48 hpi. Significant difference was analyzed from three replicates. The circles indicated lesion areas. Error bars indicate the mean fold changes ±SD (**P < 0.01, Student’s t-test). **(E-G) Plant resistance to *P. capsici* induced by RPH1 is dependent on SOBIR1**.

**Fig. S4.**
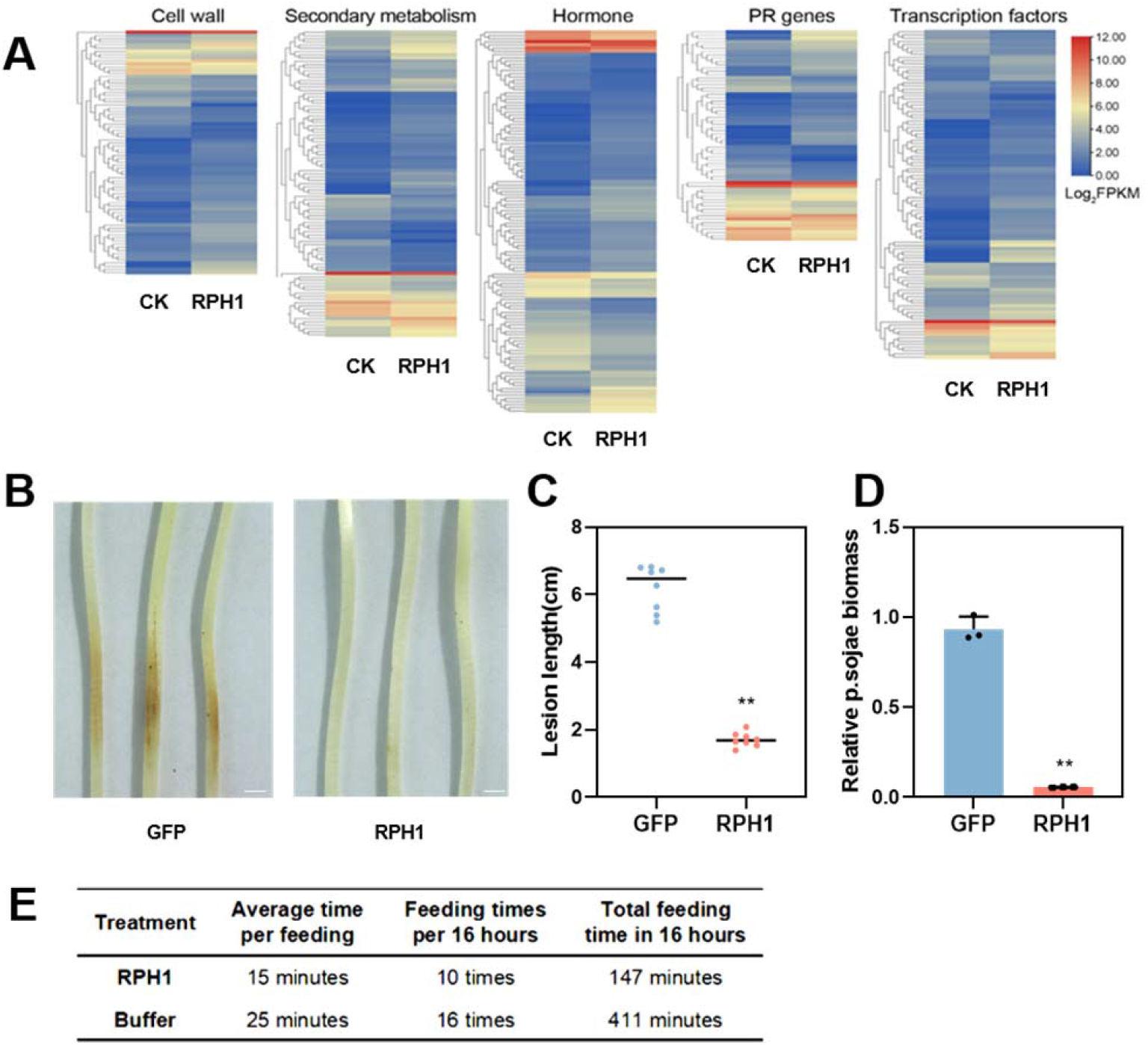
RPH1 induced PTI responses in soybean. **(A) Heat map of the expression profiles of DEGs involved in plant cell wall metabolism, secondary metabolism, hormone metabolism, pathogenesis-related genes, and transcription factors in soybean (B) RPH1 treatment protected the etiolated hypocotyls of soybean (cultivar Hefeng 47) from *Phytophthora sojae* infection**. The photographs were taken at 48 hpi and the experiments were repeated at least three times. Bar = 1 cm. **(C and D) RPH1 treatment significantly reduced the disease lesion formation and relative pathogen biomass.** Infected leaves were collected 48 hpi for genomic DNA quantitative PCR analysis. GmELF1β was used as a reference gene. Statistical differences were calculated from eight biological replicates and error bars indicate means ± SD (**P < 0.01, Student’s t-test). **(E) Analysis of feeding time differences of *R. pedestris* on RPH1- and buffer-treated soybean.**

**Fig. S5.**
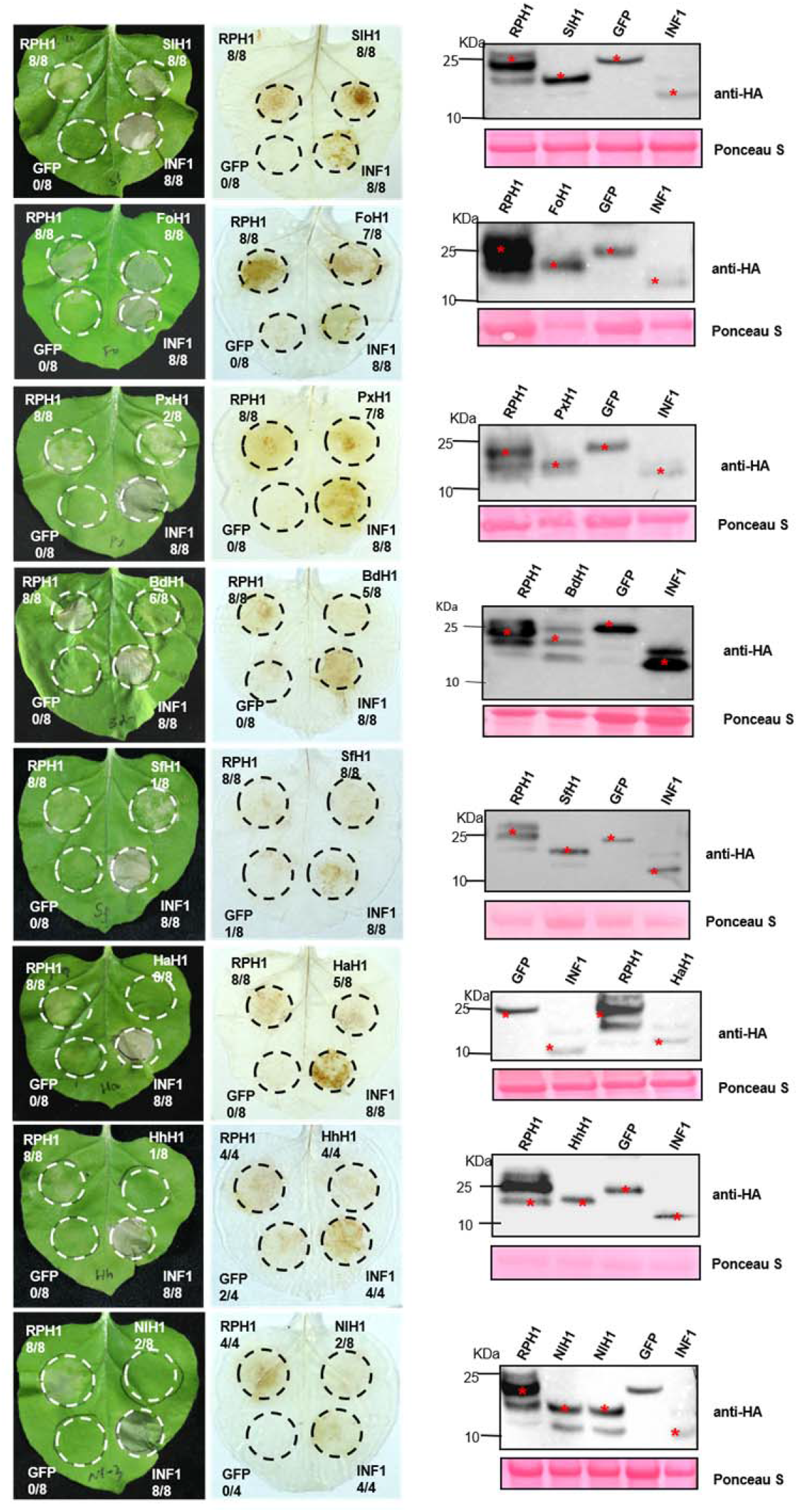
RPH1 is a conserved HAMP in herbivorous insects. RPH1 homologs from *Spodoptera litura* (SlH1), *Frankliniella occidentalis* (FoH1), *Bactrocera dorsalis* (BdH1), *Plutella xylostella* (PxH1), *Helicoverpa armigera* (HaH1), *Halyomorpha halys* (HhH1), *Nilaparvata lugens* (NlH1) were transiently expressed in *N. benthamiana*. ROS accumulation was stained by DAB. Photographs were taken 5d post infiltration. Western blot detection of proteins expressed in *N. benthamiana* using anti-HA antibodies. The red asterisks (*) indicate the predicted protein sizes. Ponceau staining of Rubisco protein (PS) indicated that equal amount of each sample was loaded.

**Fig. S6.**
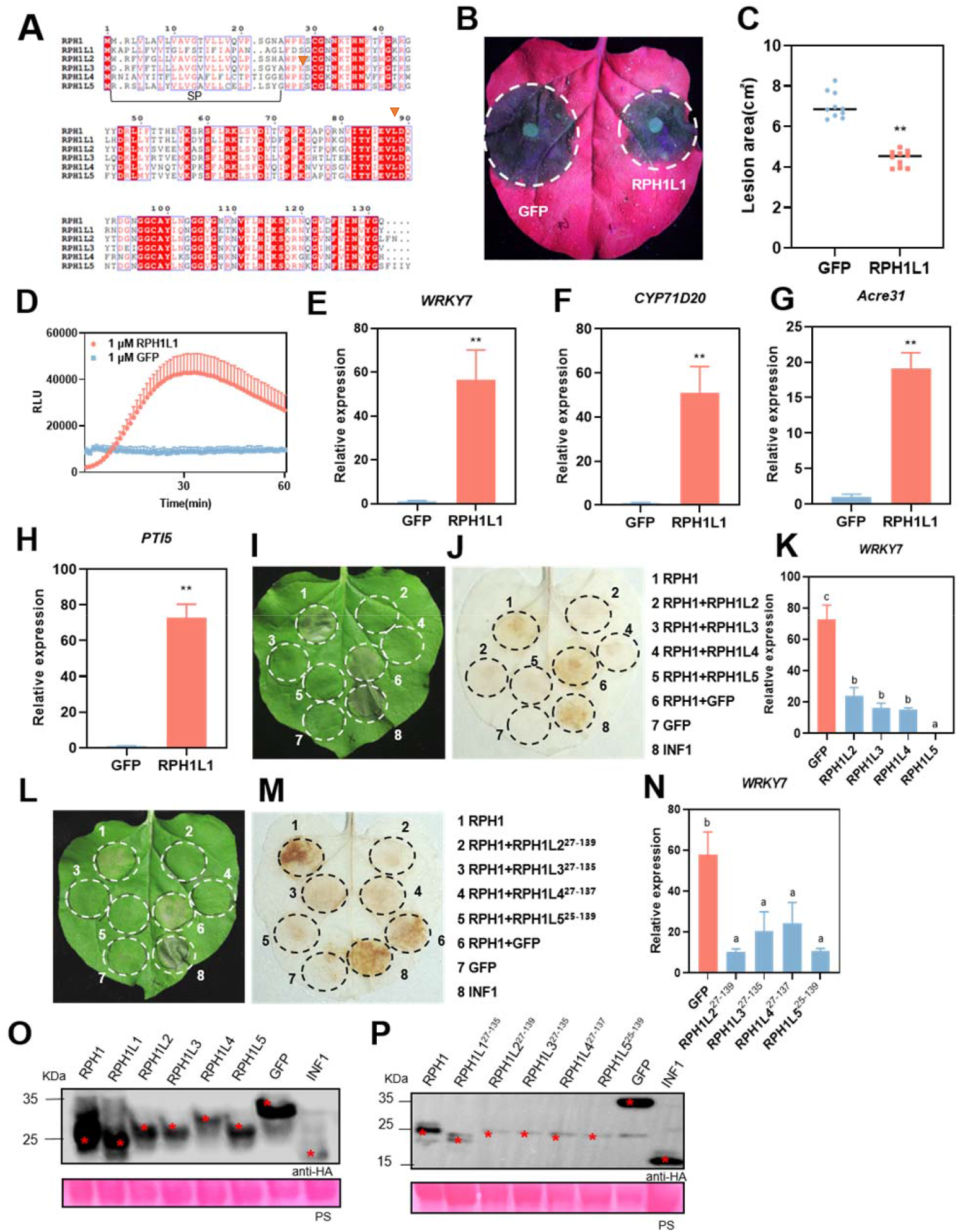
*R. pedestris* evolved paralogous genes to evade plant immunity triggered by RPH1. **(A) Protein sequence alignment of RPH1 and its paralogs using ClustalW. (B and C) RPH1L1 protein significantly increased plant resistance against *P. capsici***. Photographs were taken 48 h post inoculation (hpi) under UV light. The lesion area was measured at 48 hpi and calculated from eight biological replicates. Error bars indicate means±SD (**P < 0.01, Student’s t-test). **(D) Full length of RPH1L1 triggered ROS burst in *N. benthamiana***. GFP served as a control. ROS production was measured by the peroxidase-luminol assay. Mean RLU (Relative Luminescence Unit) (±SD) are shown (n=14). **(E-H) RPH1L1 increased the expression levels of PTI marker genes in *N. benthamiana*..** Relative expression was quantified by qRT-PCR using *Nbactin* as a reference gene. Bars indicate mean fold changes ± SD (**P < 0.01, Student’s t-test). **(I and J) Four paralog genes with SPs of RPH1 (RPH1L2, 3, 4, 5) suppressed RPH1-induced cell death and ROS accumulation in *N. benthamiana*.** GFP was used as the negative control and INF1 was the positive control. Photographs were taken 5d post infiltration. **(K) Up-regulation of *WRKY7* induced by RPH1 was inhibited by four paralog genes with SPs (RPH1L2, 3, 4, 5)**. *Nbactin* was used as an endogenous control. Means and standard errors from three independent replicates are shown. Significant differences were indicated by different letters (P < 0.01; Duncan’s multiple range test). **(L and M) Four paralog genes without SPs inhibited RPH1-induced cell death and ROS accumulation in *N. benthamiana*. (N) Increased relative expression of *WRKY7* induced by RPH1 were compromised by four paralog genes without SPs. (O and P) Western blot detection of proteins expression using anti-HA antibodies**. The red asterisks (*) indicate the predicted protein sizes in kDa. Ponceau staining of Rubisco protein (PS) indicated that equal amount of each sample was loaded.

